# Thermal Refuges as Extended Phenotypes: Lodge Construction by Bush Karoo Rats

**DOI:** 10.1101/2025.07.28.667257

**Authors:** Carsten Schradin, Nkululeko Mbanjwa, Nkululeko Nyawo, Nicola Stievenard, Perfect Dlamini, Neville Pillay, Lindelani Makuya

## Abstract

Animal architecture has received considerable attention as an extended phenotype that buffers organisms against environmental harshness. However, few studies have described animal architecture in a representative sample, assessed and then tested its function experimentally. We adopted such an approach for stick lodges built by the bush Karoo rat (*Otomys unisulcatus*), a solitary rodent inhabiting semi-arid regions of southern Africa. These lodges are among the largest animal-built structures by solitary mammals, yet their structural variation and ecological function remain poorly quantified. We conducted the first population-level survey of all lodges in our 4.5 ha field site to assess external and internal lodge structure, assess their effects on microclimate, and tested experimentally whether lodge size affects microclimate. Lodges were several hundred times larger than their builders and featured complex structure, including basking platforms, nesting, food and latrine chambers. Inside lodges, temperatures were closer to the bush Karoo rat’s thermoneutral zone and exhibited higher humidity than ambient conditions, which could reduce costs of thermoregulation and reduce water loss. These benefits correlated with initial lodge size and decreased when lodge size was experimentally reduced. Tunnels were often blocked off with thorny branches, possibly to deter snakes. Platforms typically faced eastwards and were regularly used for basking, likely enhancing passive heat gain. We conclude that bush Karoo rat lodges are complex, and costly to build and maintain. They improve thermoregulation and provide predator protection and food storage. Bush Karoo rat lodges (1) are external structures, adding to the builder’s body (2) are functional, (3) probably increasing fitness, (4) result from behavioural activity, and (5) are likely genetically encoded. We therefore consider them an extended phenotype shaped by ecological pressures and evolutionary history.

## Introduction

Many animals modify their environment by constructing nests, burrows or shelters, collectively considered as animal architecture (Laidre 2021). Social insects such as bees, wasps, ants and termites are well known for building elaborate nests and mounds, often thousands of times larger than a single individual (Hölldobler & Wilson 2009). Fish, such as sticklebacks and many cichlids, construct nests for spawning and rearing young (Stiassny & Meyer 1999; Östlund-Nilsson 2000). Most birds build nests for breeding (Perez, Manica & Medina 2023); male bower birds create elaborate colourful structures to attract mates (Marshall 1944; Marshall 2008), and male malleefowls (*Leipoa ocellata*) construct large compost heaps to incubate eggs (Stenhouse & Moseby 2022). Such examples of animal architecture are considered extended phenotypes (Dawkins 1982), shaped by genetic and developmental processes and often linked to increased fitness (Woods *et al*. 2021).

Mammals also produce a variety of animal architectures, some rivalling those of social insects and birds in complexity. Some bats build roosts of leaves, modify termite mounds or bird nests to create their own shelter (Page & Dechmann 2022). Beavers (*Castor spec*) are renowned for building extensive dams, architecture that not only provide refuge but reshape entire ecosystems (Wilson & Bremner-Harrison 2025). Nearly half of all mammal species create underground burrows (Benevento 2025), for example wombats (*Lasiorhinus latifrons*; (Carver, Stannard & Martin-Allen 2023), aardvarks (*Orycteropus afer*; (Taylor & Skinner 2003), and many carnivores (Mickevičius 2002). While muroid rodents are well known for digging burrows (Kinlaw 1999), some of them construct aboveground lodges from plant material (Qiu & Schradin 2024), such as north American pack rats (*Neotoma* spec.; Betancourt, Van Devender & Martin 1990), Australian greater stick-nest rat (*Leporillus conditor*; (Pearson *et al*. 1999), and the African acacia rat (*Thallomys spp.*) (Eccard *et al*. 2006; Meyer *et al*. 2008). While these structures are well documented across species, little is known about intraspecific variation in architectural feature.

Studying animal architecture is often very time-consuming and methodologically challenging. For example, determining the length and complexity of wombat warrens first involved a student physically entering the tunnels (Nicholson 1963). Only much later were the internal structures studied with modern camera equipment inserted through surface-drilled holes (Shimmin, Skinner & Baudinette 2002). Drilling holes is costly in time and resources, which may explain why only seven warrens per site were sampled. European badgers (*Meles meles*) are widely studied, and the location of their dens and the number of entrances has often been recorded (Macdonald *et al*. 2004), but few studies have investigated their internal structure. One exception is Roper (1992) who excavated 19 warrens and found significant variation in the number of chambers (1 to 78), latrines (0 to 8) and many other parameters. For more complex structures than tunnel systems, the situation becomes even more intricate. Middens of pack rats partly consist of a very hard concrete like crystallized urine (called amberat), hindering structural analysis (Betancourt, Van Devender & Martin 1990). While the composition of large middens has been studied to reconstruct past environments (Betancourt, Van Devender & Martin 1990; Elias 2013), their internal structure and variation remain poorly understood (Bonaccorso & Brown 1972). In general, many studies on animal architecture focused on impressive examples rather than on a representative sample.

The study of animal architecture aims to understand how organisms modify their environments in ways that influence fitness, often forming what Dawkins (1982) termed “extended phenotypes”. These structures are not just behavioral by-products, but evolved traits shaped by selection. To qualify as an extended phenotype, a structure must meet five key criteria (Dawkins 1982; Laidre 2021): i) it must be external to the organism; ii) result from its behavior; iii) provide functional benefits; iv) enhance fitness; and v) exhibit properties that reflect genetic influence. However, most studies of animal architecture focus on exceptional structures, rarely quantifying intraspecific variation or directly testing function. This hampers our understanding of the ecological pressures and evolutionary mechanisms shaping these traits. Assessing structure-function relationships in animal architecture requires an integrative approach to describe the structure, assess its consequences for the animals, and experimentally test its function (Laidre 2021).

The bush Karoo rat (*Otomys unisulcatus*) is endemic to semi-arid areas of South Africa and is known for constructing large stick lodges (Fig. 1). These lodges can reach 70cm in height, span over two meters in diameter and contain up to 13 000 sticks weighting over 8 kg (Vermeulen & Nel 1988). While Vermeulen and Nel (1988) reported mean values for 70 lodges, their sampling strategy is unknown, and only nine lodges were fully deconstructed to study the internal structure. Consequently, their reports of features, such as the presence of two nesting chambers and two latrines, in every lodge cannot be generalized, especially to smaller lodges. Other research has reported external structures not mentioned by Vermeulen and Nel (1988), such as platforms potentially used for basking (Stuart & Stuart 2019). Bush Karoo rat lodges are structurally complex, comprising of a variety of materials and components. Yet, no study to date has provided a detailed overview of the variation of lodge size and structures in a representative sample, and it remains unknown whether and to what extent specific structures such as platforms are consistently present. Therefore, the extent to which this architecture functions as an extended phenotype remains unclear.

**Fig. 1.**
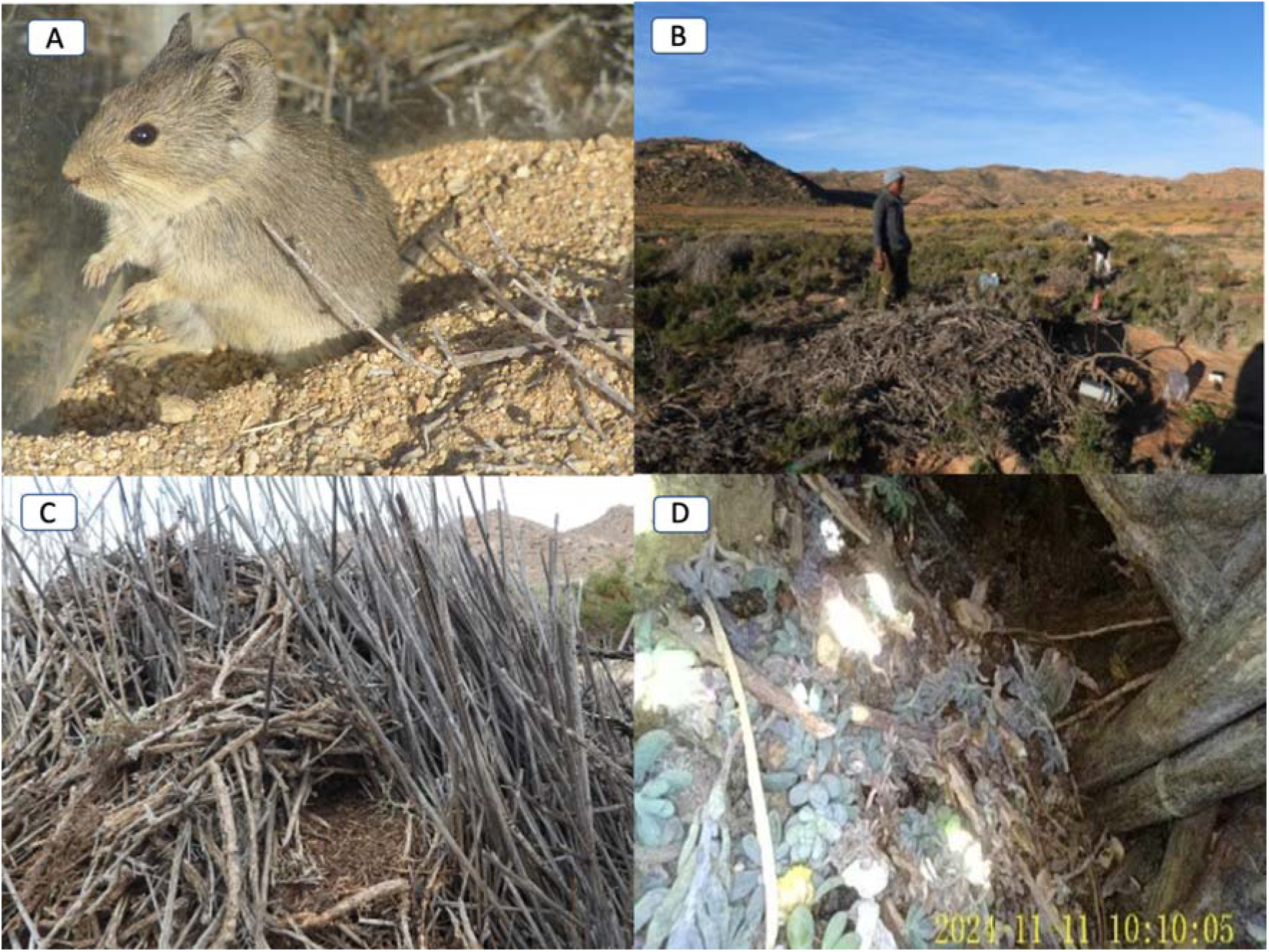
(A) The bush Karoo rat builds (B) extensive stick lodges with (C) prominent external structure such as platforms used for basking and (D) internal structure such as specialized food chambers (shown here); nest chambers and latrines are also present (nor shown here).

The architecture of Bush Karoo rat lodges likely serves multiple functions. Externally, raised platforms may facilitate sun basking, a behaviour observed in other sympatric rodents (Schradin *et al*. 2007). Internally, lodges appear to buffer extreme temperatures, providing a stable microclimate with cooler day and warmer night temperatures as well as higher humidity when compared to ambient (du Plessis, Kerley & Winter 1992; Jackson *et al*. 2002). However, the extent to which this climatic buffering depends on lodge size remains unknown, with studies differing considerably in the estimated effect of lodges on microclimate; du Plessis, Kerley and Winter (1992) reported temperature differences of 10- 11°C compared to ambient conditions, but Jackson *et al*. (2002) reported a difference of only 2° C. Experimental approaches to understand the function of bush Karoo rat lodges are so far missing.

Our goal was to test whether bush Karoo rat lodges function as extended phenotypes. For this, we studied the external and internal structure of bush Karoo rat lodges in a representative sample to better understand the function of this animal architecture, capturing the full spectrum of lodge size variation. We quantified architectural features, including size, entrances, platforms, tunnels and construction materials used, and assessed their relationship to environmental conditions and behaviour. We predicted that lodge height would correlate with other structural traits, indicating coordinated construction. We investigated whether platforms exist and if so, whether bush Karoo rats would use them for basking. If platforms are especially built for basking, we expected them to face east or north-east, so they are exposed to sun during cooler morning. We assessed how lodge size varied seasonally, either increasing during the dry season due to building activity or decreasing due to desiccation of moist building material added in the moist season (Sangweni *et al*. 2025). Using an endoscope camera, we explored internal structures and identified specialized chambers. We measured internal lodge temperatures and humidity to test whether (a) lodges buffer against environmental extremes, and (b) larger lodges provide stronger microclimatic benefits. To test this experimentally, we deconstructed eight unoccupied lodges and predicted that the microclimate will be less favorable when lodges are deconstructed by 50% and by 75%, directly testing for an effect of lodge size on function as extended phenotype. Our integrative approach allowed us to test the ecological function of one of the largest examples of solitary mammal architecture and address whether the lodges qualified as an extended phenotype shaped by natural selection.

## Material and Methods

### Study Species

The 100g heavy diurnal bush Karoo rat (Fig. 1) is endemic to the semiarid Karoo and the Succulent Karoo of South Africa (Do Linh San *et al*. 2016). It constructs stick-lodges inside shrubs (Fig. 1), which provide a favourable microclimate with high humidity and mild temperatures for protection against the external environment (Brown & Willan 1991; du Plessis & Kerley 1991). Stick lodges on our field site can be more than 20 years old and are used by consecutive generations over many years (Schradin & Makuya, unpubl. data). The bush Karoo rat is a central-place forager, foraging within less than 25m from its lodge and before returning to it (Makuya, Pillay & Schradin 2024).

### Study Site and Study Period

We conducted our study on our 4.5 ha site used for long-term studies on bush Karoo rats since 2018. It is adjacent to the Succulent Karoo Research Station in the Goegap Nature Reserve, Northern Cape, South Africa from August 2024 to April 2025. The dominant vegetation includes *Tetraena retrofracta* and *Lycium cinerum* shrubs interspersed with sandy areas which in spring are covered by succulents such as *Mesembryanthemum guerichianum* and ephemerals (Rösch 2001). The Succulent Karoo experiences a cold moist season in winter and spring, and a hot dry season in summer and autumn. Our field site receives an average annual precipitation of 160mm, mainly in winter, when frost is common (minimum night temperature -3°C), with day temperature always being above zero and ranging from 10- 25°C (data from our weather station at the research station). In summer, day temperatures are typically above 30°C and can reach 40°C, with night temperatures being between 5°C and 25°C.

### Determining external lodge structure

We recorded external structural measurements of all 106 occupied lodges on the field site during the moist season (September–November 2024). During the dry season (January– February 2025), we surveyed all 141 occupied lodges, including 97 that remained occupied from the moist season, 34 older lodges that were previously unoccupied, and 10 newly constructed lodges.

For every lodge, we recorded its unique ID and the ID of the bush Karoo rat occupying it, based on trapping data (for details of trapping see (Makuya, Pillay & Schradin 2024). The shrub species into which the lodge was built was recorded. Lodge height (cm) was measured using two 180 cm wooden rulers: One ruler was placed vertically from the ground, and a second wooden plank ruler was placed horizontally at the top of the lodge to meet the first wooden ruler. The diameter of the lodge was measured with the same wooden ruler. A 30 m tape was used to measure the lodge’s circumference. We estimated the percentage composition of building materials, including dry sticks, rodent faeces, ungulate dung (from mountain zebra, gemsbok and springbok), bones (from large mammals), clay and stones. The number of stones and bones were also quantified. We counted the total number of entrances per lodge.

*Identifying and measuring lodge platforms:* Platforms (Fig. 1C) were defined as specialized parts of the external lodge structure which are horizontally flat surfaces on which a bush Karoo rat can sit or lie down to sun-bask or feed. We recorded the direction the platform was facing as cardinal directions in degrees using a phone compass. We classified platforms according to their exposure to the sun as open, half-open or closed, and whether each platform contained an entrance. The longest horizontal side of the platform was measured and recorded as its diameter and we measured how many cm above ground the platform was.

### Basking behaviour

We collected behavioural data from late August 2024 till April 2025 through direct behavioural observations of occupied lodges. Before each observation session, we counted the number of platforms per lodge and recorded their direction from the centre of the lodge using a compass application on a smart phone. Observations lasted for 30 minutes and started 5 minutes before sunrise or 25 minutes before sunset. Bush Karoo rats at our site are habituated to humans. Observations were conducted from a distance of 5-10m using binoculars. We collected data using animal focal sampling and one zero recording at 1- minute intervals. Basking was defined as an individual remaining stationary while exposing its body to direct sunlight. In each occurrence, we recorded the location where basking occurred: on a platform, on the lodge or on the ground.

### Examining internal lodge structure

We examined the internal structure of lodges from the 28^th^ September to the 11^th^ December 2024, spanning the moist to the onset of the dry season. The same 106 lodges used for external structure during the moist season were included. We used a Kentfaith AGC-500L flexible fibre-optic dual endoscope camera (front and side cameras) with high-illumination white LED for imaging, 5inch colour IPS screen and 1080P resolution. The camera was mounted on a 3m, 8.5 mm diameter cable, marked with colour-coded tape at 10cm intervals. The endoscope camera was bidirectional at 180°, allowing for 90° movement to both sides.

To measure tunnel length, the marked cable was inserted through each entrance. The length was then measured using the markings together with a ruler. From the video footage, we recorded any food or other material inside the tunnels, and whether it led to any chamber, or to the outside, or were blocked by sticks that made it impossible to insert the cable any further. If tunnels lead to chambers, we recorded whether the chambers contained faeces, food, nesting material or were empty. If a nest was found, we photographed it but did not insert the camera further to avoid disturbing the nesting site.

### Lodge deconstruction

From February to April 2025 (dry season), we deconstructed eight unoccupied lodges, based on trapping records, absence of fresh faeces or food remains, and lack of active runways. In addition, each lodge was also observed for at least two 30-minute sessions during early morning or late afternoon activity peaks to confirm lack of occupation. Three lodges were located outside the field site and we did not know their occupancy histories. The remaining five lodges had not been occupied within the previous year.

An iron rod (5mm diameter), marked with tape every 10cm, was inserted into each lodge until it reached the ground, but without it sticking into the ground. Lodges were deconstructed in 10cm layers (except the shorter topmost layer), with sticks removed individually. For each layer, sticks were sorted into five lengths categories: 1-5cm; 5-10cm; 10-15cm; 15-20cm; and >20cm. We recorded the number and total mass (to nearest gram) of sticks per category. Additionally, we measured the thickness (to the nearest 0.5 mm) of 12 randomly chosen sticks per category using a Vernier scale. We also recorded the mass of any other materials, such as dung, bones, stones, and anthropogenic debris (e.g. wires, metal pieces, glass), and the mix of clay/faeces/sticks smaller than 1cm.

*Internal structure.* We sketched internal features, focusing on whether branches served as anchor points or structural supports. Plastic pipes (3.5 cm diameter) were inserted into visible tunnels to preserve their shape during deconstruction and measured tunnel lengths. The intention was to assess, during deconstruction, whether the tunnel continued beyond the pipe, perhaps forming a curve, but this method proved ineffective: in all cases, the space beyond the pipe was obstructed by sticks.

When we encountered soft material, sticks were removed carefully to identify potential nests, before removing them from the lodge. We recorded nest dimensions and mass, but could not measure nesting chamber size, because chambers collapsed during deconstruction. No latrine chambers were found, also likely due to structural collapse. The only recurring internal feature was a “ceiling” structure in the bottom layers, composed of small sticks, sand, clay and faeces.

*Temperatures and humidity measures:* We measured internal temperature and relative humidity using Hygrochron DS1923 iButtons (8KB memory; Maxim Integrated). In each of the 8 lodges selected for deconstruction, one iButton was inserted into a tunnel (using 60cm long tweezers) at least 24 hours before deconstruction began. As control, a second logger was placed on the ground in the shade under a shrub nearby that was of the same species and similar size as the lodge shrub. All iButtons were enclosed in an iButton holder, which was attached to a thin wire and secured into the ground to prevent removal by animals.

Deconstruction was paused for at least 24 hours when each lodge reached 50% and 75% of its original height. This allowed for 24 hours recordings before deconstruction, after 50% and after 75% of deconstruction.

Loggers were programmed to collect data every 5 minutes. Limited memory capacity on the iButtons led to data loss in the first deconstructed lodge (which took longer to deconstruct), reducing the usable sample to 7 lodges. In another lodge, internal logger readings exceeded 50°C after 75% deconstruction, likely due to direct sun exposure, only data from before deconstruction and after 50% deconstruction were used. One logger consistently recorded relative humidity around 100% (range 95–110%) with minimal variation, way above measures from other lodges and ambient (Fig. 7). We considered this sensor faulty and excluded its data. Heavy rainfall during one deconstruction led to elevated humidity levels, making the data from the lodge unsuitable for evaluating humidity readings. For humidity, the final sample sizes were therefore reduced to N=5 lodges (N=6 for before deconstruction data).

### Statistical Analysis

All statistical analyses were conducted in R (version 4.3.1; R Core Team 2022). For correlations we used Pearson’s r. To assess the relationships between different measures of external lodge characteristics, we used linear mixed model (LMM; package lme4 in R) with lodge height as dependent factor and shrub species and season as co-variates and ID of the lodge as random factor. Our aim was not to find the best model explain lodge height, but to see how it correlates with other lodge measurements while controlling for shrub species and testing for a seasonal effect. Thus, we run four different models, each of which either had diameter, circumference, number of entrances or number of platforms as continuous variable. A paired t-test was used to compare lodge characteristics (i.e. lodge height, diameter, circumference, number of entrances, and number of platforms) during the moist and dry seasons for lodges present in both seasons.

To test whether platform orientations deviated from uniformity, we used dry season data and applied the Hermans–Rasson test, a robust and powerful tool for analysing circular distribution of compass data (Landler et al., 2019). We used data from the dry season, when more lodges were measured. Tests were run first with all platforms followed by only open platforms that would facilitate basking To assess whether bush Karoo rats used platforms for basking, we analysed only those observation sessions where at least one platform was present and the focal rat was observed basking, resulting in 125 observations (91 morning and 34 in the afternoon) of 55 different individuals from 57 different lodges. For each individual, we calculated the percentages of basking events occurring on platforms, elsewhere on the lodge, or on the ground. The data were not normally distributed, so we used the non-parametric Friedman test for paired data followed by post hoc comparisons.

To assess whether larger lodges were associated with greater microclimate buffering, we ran Pearson’s correlations between lodge height and the absolute difference between internal and ambient temperature and humidity prior to deconstruction.

A favourable microclimate is achieved when lodge temperatures are closer to the thermoneutral zone (TNZ) of an animals than ambient temperatures. Therefore, we calculated the temperature deviation from the thermoneutral zone of bush Karoo rats, which is between 24°C and 30°C (du Plessis, T. Erasmus & Kerley 1989). For each value, those within the TNZ were scored as zero and values outside of the TNZ were assigned the absolute distance to the nearest TNZ boundary.

We ran two linear mixed models to test whether lodge temperatures differed from ambient temperatures depending on deconstruction level, using data from the hottest hours of the day (10-18h; Figure 6). To assess whether lodges buffered against cold, we used data from the coldest part of the day (20-8h, Figure 6).

We then modelized a LMM to see the effect of the deconstruction level on the absolute value of the deviation from TNZ. To analyse the humidity data, we fitted a LMM to analyse the effect of the level of deconstruction on lodge humidity compared to ambient humidity. For all models, we included 3 random effects: lodge identity, to account for repeated measures within lodges and for the interindividual variation between lodges not related to size. Hour of the day because temperatures every hour are highly correlated due to the circadian cycle. Date to account for the effect of seasonality because the ambient temperature decreases during late summer and autumn and affect the data.

All data and code will be published after acceptance (Schradin *et al*. 2025) and are available for review here: https://data.indores.fr:443/privateurl.xhtml?token=c3bb7013-9a3a-4d69-8f0e-0381c01401da

### Text editing

All text was first written and edited by us. Selected sections of the introduction and discussion were refined in ChatGPT using the prompt “Act as a scientific editor; Change the text below to read better; Also correct any grammatical errors; Keep it short and precise; Suggest all changes and mark them for me to review.” The suggested changes were then reviewed and incorporated only when they improved the manuscript. The new version was then still read and edited by three of the authors (CS, NP and LM).

### Ethical Note

The research was conducted under a national South African section 20 research permit and trapping and observation was approved by the animal research ethics committee of the University of the Witwatersrand (AREC23/08/005).

## Results

### External lodge structure

Lodges varied considerably in size (Table 1) and were constructed mainly of sticks, incorporating other natural or anthropogenic materials (Table 2). Of the 106 lodges recorded in the moist season, 82 (77%) were built in *Tetraena retrofracta* and 18 (17%) in *Lycium cinerum* shrubs, with the remaining six distributed in other shrub species. The two shrub species differ in their growth characteristics, with *T. retrofracta* shrubs forming broad canopies but *L. cinerum* shrubs growing taller and upright. To test whether shrub characteristics influenced lodge size, we compared lodges constructed in these species during the moist season, when all lodges were several years old (excluding the dry season when some were newly built). Lodges in *T. retrofracta* (N = 82) did not have a significantly larger diameter (p = 0.69, t = 0.40) but were significantly shorter than those in *L. cinerum* (N = 18; p < 0.0001, t = 4.311; Fig. 2).

**Fig. 2.**
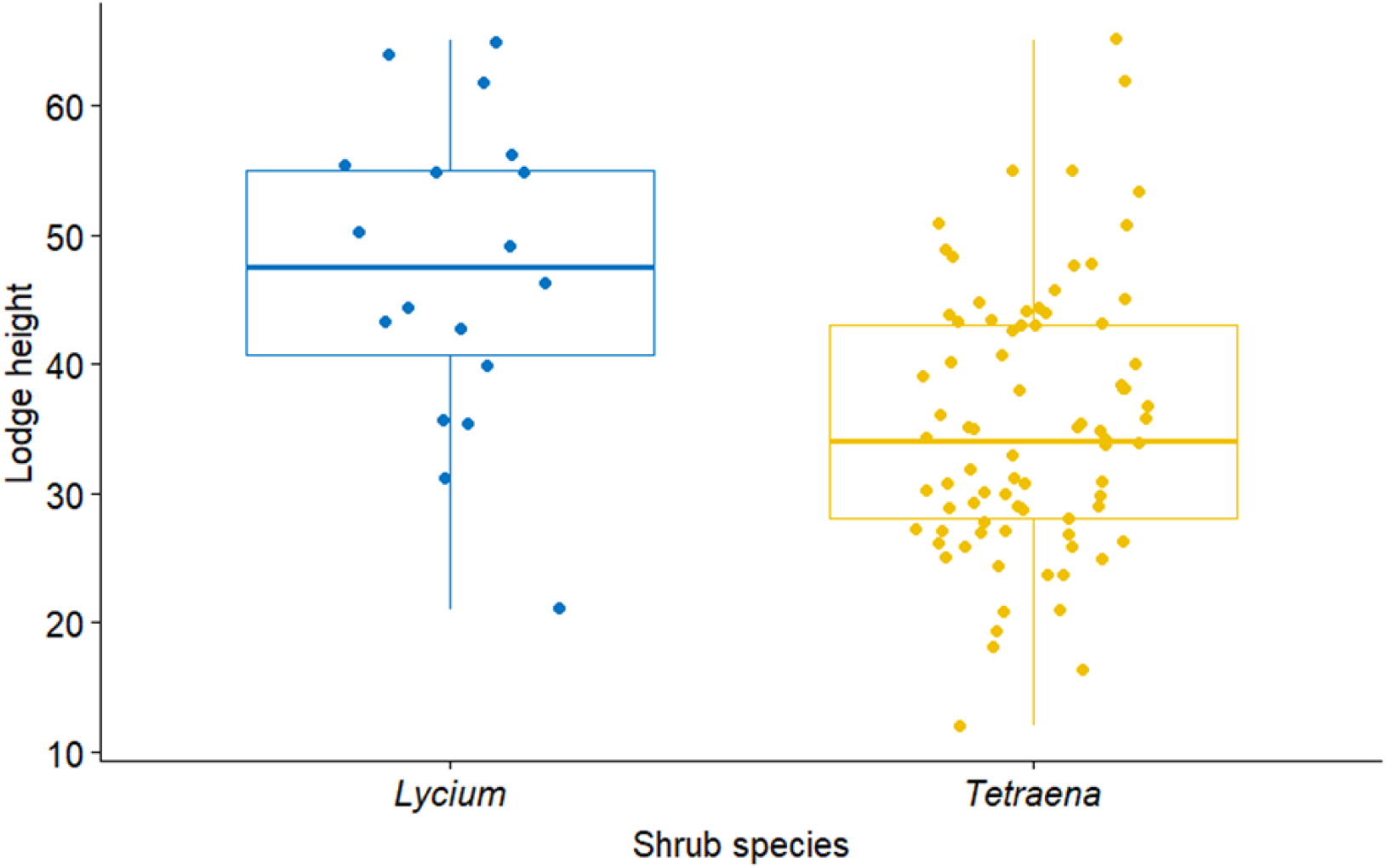
Height of lodges depending on into which shrub species they were constructed in, the high growing *Lycium cinerum* or the broad growing *Tetraena retrofracta*. Boxplots show 1st and 3rd quartiles, the horizontal line indicates the median, the whiskers represent the minimum and maximum of the outlier data and points represent individual values.

**Table 1.** Descriptive statistics of external lodge characteristics, expressed as mean + SD (range)

|  | Moist season (N=106) | Dry season (N=141) |
| --- | --- | --- |
| Circumference | 366 ± 111 cm (154 – 1102cm) | 325 ± 137 cm (58- 1318 cm) |
| Diameter | 113 ± 34 cm (30 – 240 cm) | 110 ± 37 cm (23 – 275 cm) |
| Height | 38 ± 12cm (12 – 65cm) | 39 ± 12 cm (13 – 74 cm) |
| Number entrances | 2 ± 2 (0 – 10) | 2 ± 1 (0 - 6) |
| Number platforms | 2 ± 2 (0 – 9) | 2 ± 2 (0 - 12) |

**Table 2.**
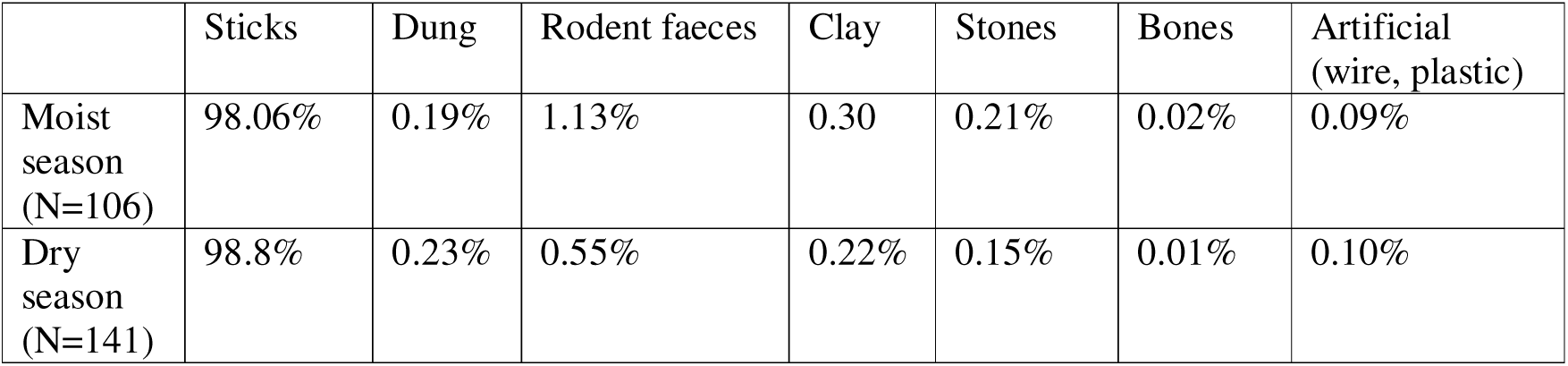
Estimates (in percentage) of different materials of which the external structure of lodges consisted.

Among the 97 lodges surveyed in both seasons, paired comparisons revealed that lodge height increased from the moist to the dry season (t = –2.878, p < 0.01; Fig. 3A), while circumference decreased (t = 5.737, p < 0.0001; Fig. 3B). Lodge diameter remained unchanged (t = –0.091, p = 0.93; Fig. 3C). Lodges had significantly more entrances in the moist season (t = 2.070, p = 0.04; Fig. 3D), while the number of platforms did not differ between seasons (t = –1.734, p = 0.09; Fig. 3E).

**Fig. 3.**
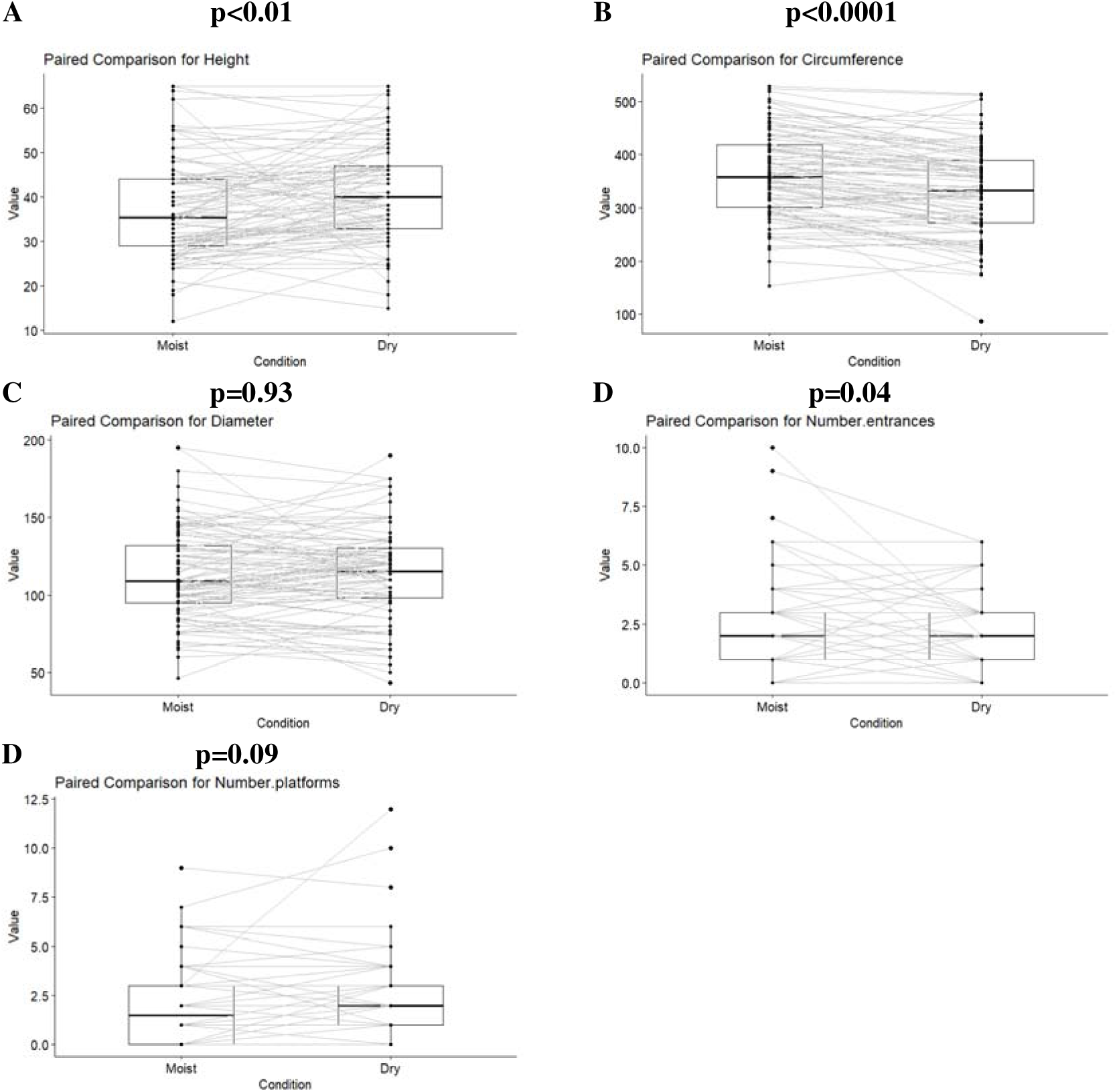
Seasonal differences in external lodge characteristics. Data of 97 lodges that were measured in the moist and dry season. The boxplots represent the interquartile range, with the horizontal line indicating the median. Whiskers extend to the minimum and maximum values, excluding outliers which are shown as individual points. Values of individual lodges are connected by a line. A: Height. B: Circumference. C: Diameter. D: Number of entrances. E: Number of platforms.

Linear mixed models with lodge species included as co-variate (see Electronic Supplement) showed that lodge height correlated positively with diameter (p < 0.0001), circumference (p < 0.0001), number of entrances (p < 0.01), and number of platforms (p < 0.001), after controlling for shrub species and season. Season had a significant effect in all models.

### Platform size and orientation

Platforms were 18.2 ± 10.5cm above ground and had a diameter of 18.0 ± 6.4cm. Most platforms were half open, protected from above but allowing sunshine in the morning/ afternoon (46% of platforms), some were closed (33%) and 21% were fully open. Platform orientation showed a significant deviation from uniformity (Hermans–Rasson test statistic = 0.0135, *p* < 0.01. N=260, Fig 4A). This pattern remained when only open platforms were considered (Hermans–Rasson test statistic = 0.0414, *p* = 0.047, N=58, Fig. 4B).

**Fig. 4.**
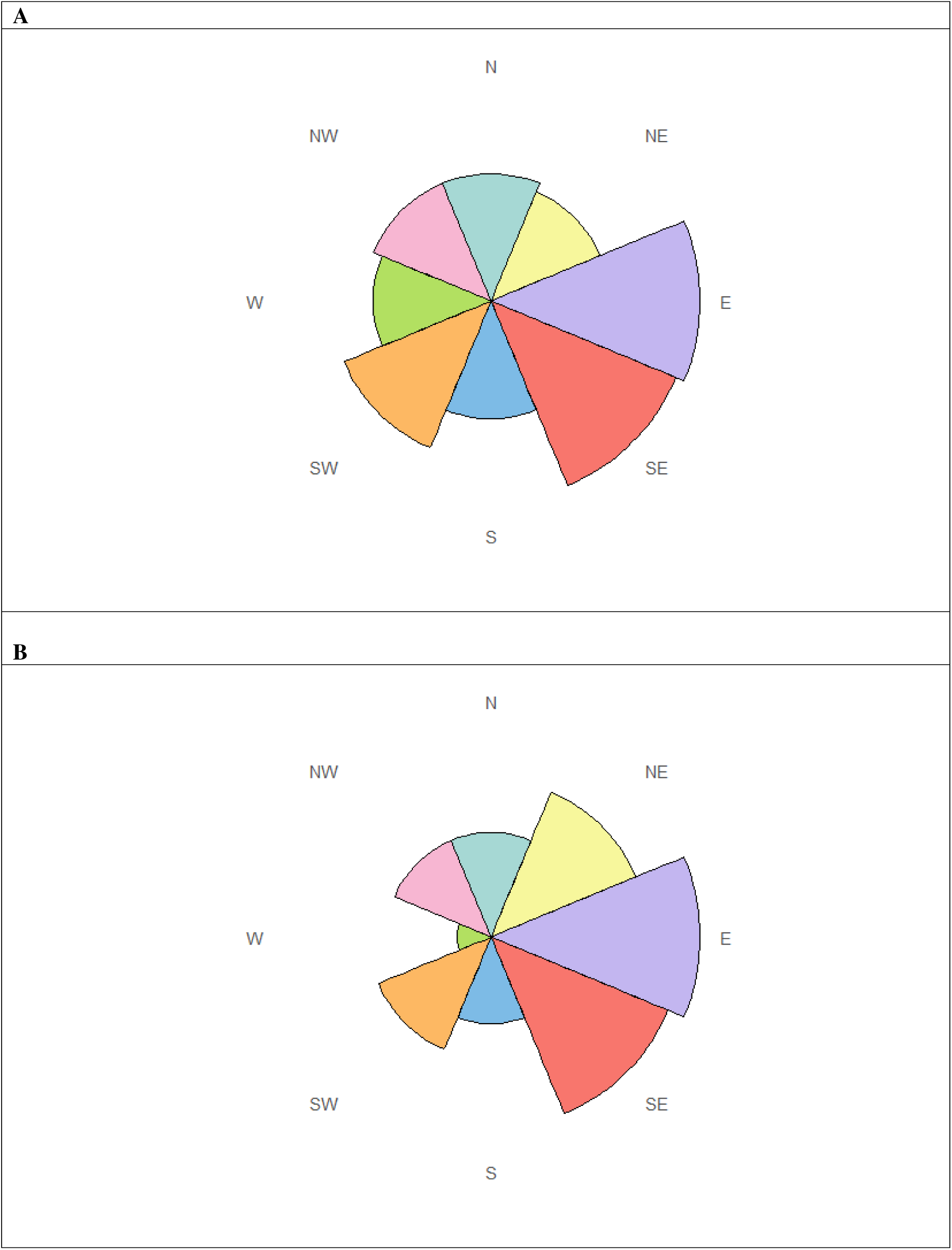
Orientation of platforms on lodges. A: All platforms. B: Only open platforms. N: north; NE: northeast; E: east; SE: southeast; S: south; SW: southwest; W: west; NW: northwest.

### Basking behaviour

Bush Karoo rats displayed significant variation in the location of basking events (Friedman test: Fr = 8.755, p = 0.01). Basking occurred most frequently on platforms (42.4 ± 43.4%), followed by the ground in front of the lodge (40.4 ± 42.3%) and least frequently elsewhere on the lodge structure (17.3 ± 31.5%). Post hoc tests showed that basking occurred significantly more often on platforms than elsewhere on the lodge (p < 0.05); other comparisons were non-significant.

### Internal lodge structure

Of the 106 lodges examined, 17 were inaccessible for the use of our endoscope camera due to dense thorny vegetation. From the remaining 89 lodges, we recorded 272 tunnels. Of these, 68 were obstructed by thorny sticks (N = 61) or clay (N=7); while 11 tunnels ended in the ground beneath the lodge; 44 tunnels lead to a chamber; and 31 tunnels lead outside. Tunnels were 34.1 ± 13.8cm long, ranging from 10cm to 75cm long. In tunnels of 10 lodges, we saw invertebrates, mainly ants, cockroaches and spiders. A total of 44 chambers were found in 73 lodges: 18 were identified as sleeping chambers, 13 contained food, 12 were categorised as latrines, and one contained both faeces and food. In only 17 lodges were we able to find a sleeping chamber, and in only one of these lodges did we find two sleeping chambers. Table S1 provides a detailed description of the internal structure, and figures S1-S5 examples of photos taken of different structures inside lodges.

### Lodge deconstruction

All eight deconstructed lodges were constructed within shrub branches, which appeared to provide mechanical support, although there was no evidence of deliberate structural shaping or branch manipulation.

In six lodges, we identified a compacted “ceiling” layer near the base, of a hard concrete like mixture of faeces, clay and sand. These concrete-like layers were found in layer 1 of all six lodges, in layer 2 of four, and in layer 3 of one lodge. Ceiling layers weighed between 1.5 and 17.5 kg (mean = 7.3 ± 6.1 kg; Table 3).

**Table 3.** Characteristics of construction material extracted from eight deconstructed lodges, expressed as mean + SD and (range for rare materials).

| Building material | Number | Mass (g) | Thickness (mm; from 12 randomly chosen sticks per size category per layer per lodge) |
| --- | --- | --- | --- |
| Sticks 1-5cm | 7764±529 | 1403 ± 1306 | 2 ± 2 |
| Sticks 6-10cm | 5065 ± 3050 | 2321 ± 2021 | 3 ± 2 |
| Sticks 11-15cm | 1195 ± 521 | 1781 ± 1655 | 4 ± 3 |
| Sticks 16-20cm | 347 ± 143 | 949 ± 931 | 5 ± 4 |
| Sticks >20cm | 167 ± 85 | 977 ± 1010 | 6 ± 5 |
| Total sticks | 14494 ± 8909 | 15023 ± 12433 | 4 ± 3 |
| Stones | | 58 ± 86 (0-229) | |
| Large ungulates dung | | 29 ± 70 (0-201) | |
| Jackal dung | | 3 ± 7 (0-19) | |
| Mix of clay, faeces and sand | | 10095 ± 10644 | |
| **Total mass of lodge material** |  | **25227 ± 21738** |  |

The internal structure of the lodges was unstable during deconstruction, making it impossible to examine tunnels and chambers. Plastic pipes inserted prior to deconstruction were obstructed, and no tunnel trajectories could be traced after the pipes. No latrines were observed during deconstruction, but 12 nests were recovered from seven lodges (1.5 ± 0.9 nests per lodge, range 0-3) Nests weighed 24.2 to 173g and were composed of soft plant material, mostly grass. Seven nests were centrally located and five were positioned laterally.

Table 3 summarises the building material that was removed during decomposition with more details per layer given in Table S2. Lodges contained a mean of 14 539 ± 8 384 sticks (range of 4 857 to 2 309 sticks). Total building mass ranged from 5.2 to 48.2 kg (mean = 25.6 ± 21.1 kg), with the smallest lodge weighing 5.2kg and the largest being 48.2kg. The size measured in height and diameter correlated with the mass of building material (r_Pearson_ = 0.81; t_6_= 3.4; p = 0.01; r_Pearson_ = 0.80; t_6_ = 3.3; p = 0.02 respectively) and the number of sticks (r_Pearson_ = 0.93; t_6_= 6.3; p < 0.001; r_Pearson_ = 0.82; t_6_ = 3.6; p = 0.01 respectively).

Compared to lodges, platforms were constructed from distinct mix material, dominated by clay and faeces (Table 4). The ratio of the mass of the building material sticks vs. other materials differed significantly between the lodges and platforms (Fisher’s exact test, χ² =58.773; p<0.0001).

**Table 4.**
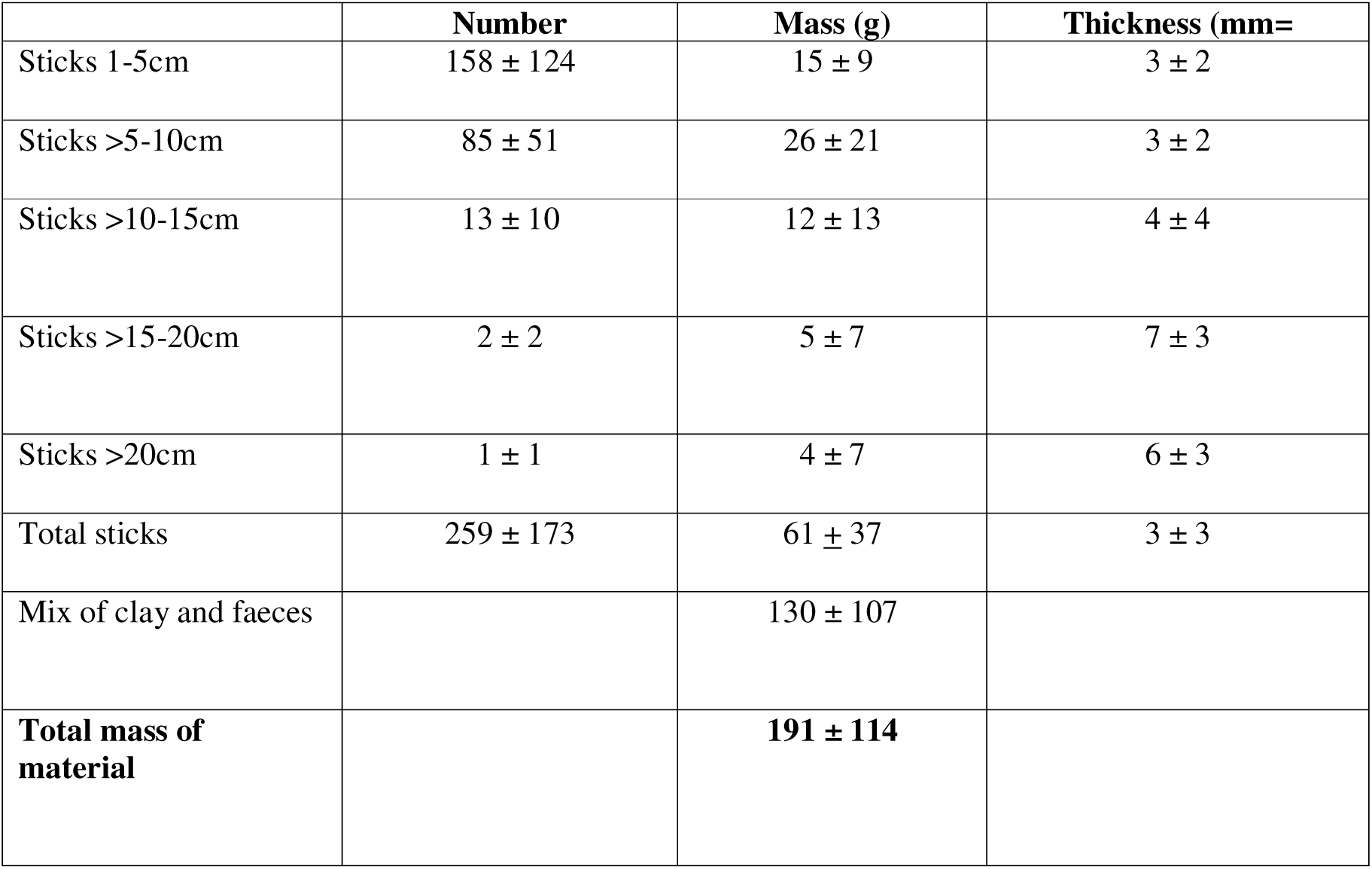
Characteristics of construction material that was extracted from six deconstructed platforms from a total of four lodges, provided as mean ± SD.

|  | Number | Mass (g) | Thickness (mm= |
| --- | --- | --- | --- |
| Sticks 1-5cm | 158 ± 124 | 15 ± 9 | 3 ± 2 |
| Sticks >5-10cm | 85 ± 51 | 26 ± 21 | 3 ± 2 |
| Sticks >10-15cm | 13 ± 10 | 12 ± 13 | 4 ± 4 |
| Sticks >15-20cm | 2 ± 2 | 5 ± 7 | 7 ± 3 |
| Sticks >20cm | 1 ± 1 | 4 ± 7 | 6 ± 3 |
| Total sticks | 259 ± 173 | 61 ± 37 | 3 ± 3 |
| Mix of clay and faeces | | 130 ± 107 | |
| **Total mass of material** |  | **191 ± 114** |  |

**Table 4:** Results of the linear mixed-effects model to identify whether deconstruction influenced lodge temperature / humidity and if it differs from ambient temperature during the hottest hours of the day (10h to 18h) and during the coldest hours of the day (20h to 7h). Significant p values are indicated in bold.

| **Model for hot temperatures** | Estimate | SE | t value | *P* value |
| --- | --- | --- | --- | --- |
| Intercept | 27.386 | 7.297 | 12.72 | **< 0.001** |
| 50% deconstruction | 0.48 | 0.20 | 2.44 | **0.015** |
| 75% deconstruction | 0.74 | 0.21 | 3.60 | **<0.001** |
| Ambient temperature | 2.23 | 0.14 | 15.95 | **<0.001** |
| **Model for cold temperatures** |  |  |  |  |
| Intercept | 19.08 | 1.33 | 14.40 | **<0.001** |
| 50% deconstruction | 0.49 | 0.11 | 4.35 | **<0.001** |
| 75% deconstruction | -0.08 | 0.13 | -0.61 | 0.54 |
| Ambient temperature | -1.92 | 0.08 | -22.69 | **<0.001** |
| **Model for relative humidity** |  |  |  |  |
| Intercept | 52.86 | 7.22 | 7.33 | **<0.001** |
| 50% deconstruction | -1.72 | 0.70 | -2.47 | **0.01** |
| 75% deconstruction | -2.84 | 0.72 | -3.94 | **<0.001** |
| Ambient humidity | -3.74 | 0.48 | -7.73 | **<0.001** |

### Lodge temperatures

Prior to deconstruction, there was a significant correlation between height of the lodge and temperature differences between lodge and ambient temperatures (r_Pearson_ = 0.76; t_5_= 2.6; p = 0.04). Intact lodges buffered temperature more effectively than those partially deconstructed, with internal temperatures deviating less from the thermoneutral zone compared to ambient conditions (Fig. 6A; Table 4).

**Fig. 6.**
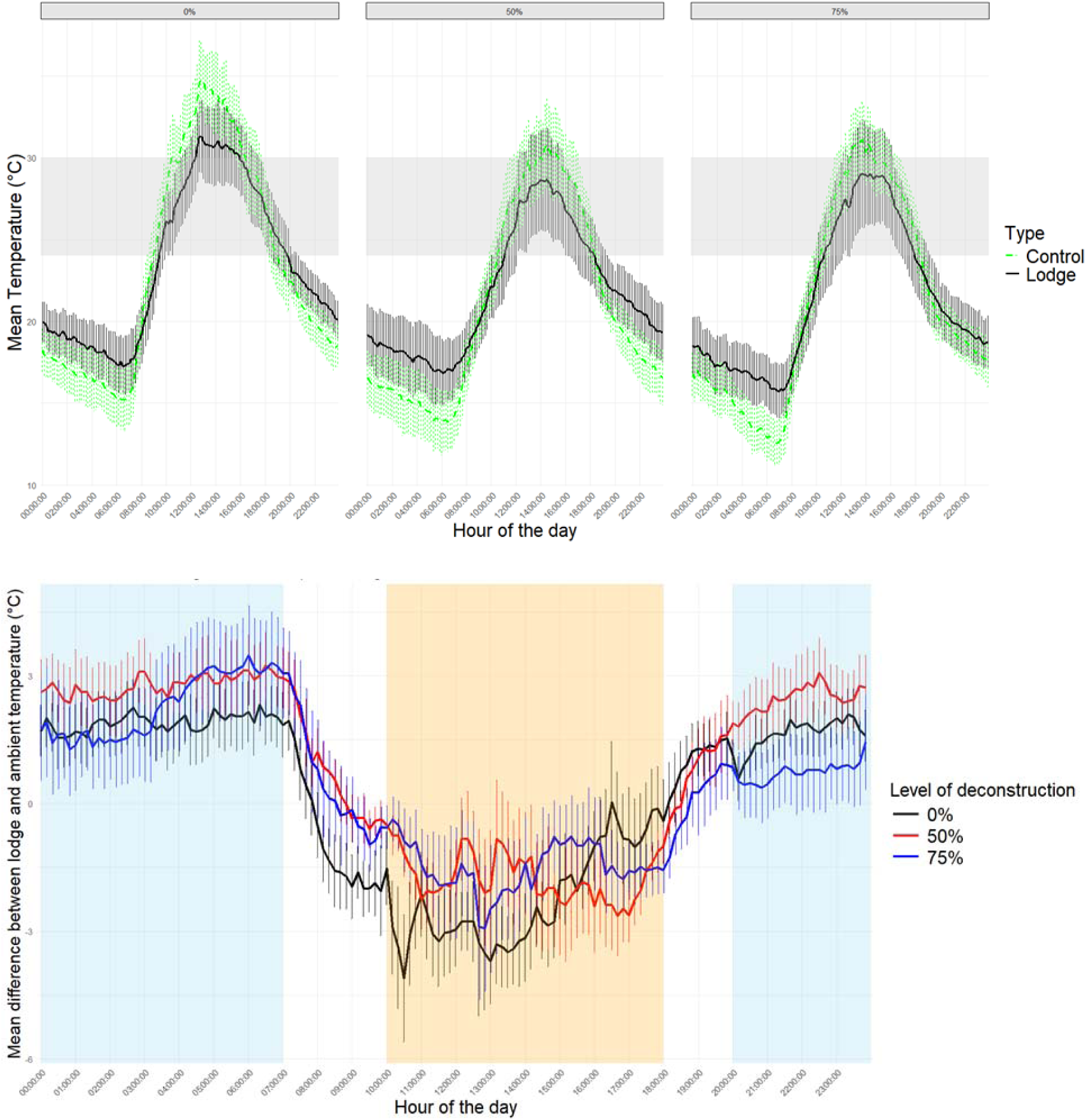
Top: Temperatures inside lodges depending on level of deconstruction and ambient temperature. Grey area: Thermoneutral zone of bush Karoo rats. Bottom: Deviation of inside lodges temperatures from ambient temperature depending on level of deconstruction. Orange shading represents the hot periods of the day, blue shading represents the cold periods, both derived from the ambient temperatures shown in the top figure. 0%: Original size. 50%: Lodge size experimentally decreased by 50%. 75%: Lodge size experimentally decreased by 75%.

During periods when it was cold, i.e., from 20:00 in the evening until 8:00 in the morning, temperatures inside the lodges were significantly warmer than ambient temperatures (p<0.0001; Table 4, Fig. 6B). There was no difference between temperatures inside intact lodges and lodges deconstructed by 75%. Surprisingly, temperatures inside the 50% deconstructed lodges were higher than in the original lodges when controlling for ambient temperatures (p< 0.0001).

During hot periods, i.e., from 10:00 in the morning until 18:00 in the evening, temperatures inside the lodges were significantly cooler than ambient temperatures (p<0.0001; Table 4, Fig. 6B). This depended on the level of deconstruction, with lodge temperatures being lowest in non-deconstructed lodges and highest in lodges deconstructed by 75% (Table 4).

The sample size for humidity in lodges before deconstruction was low due to technical problems (see methods); the correlation between lodge height and humidity was r_Pearson_ = 0.51 (t_4_= 1.2; p = 0.3). Nonetheless, relative humidity was higher in intact lodges than in ambient air (t_4594_=-7.73, p<0.0001) and in lodges deconstructed by 50% (t_4591_=-2.470, p<0.01) and lodges deconstructed by 75% (t_4583_=-3.942, p<0.0001; Table 4, Fig. 7).

**Fig. 7.**
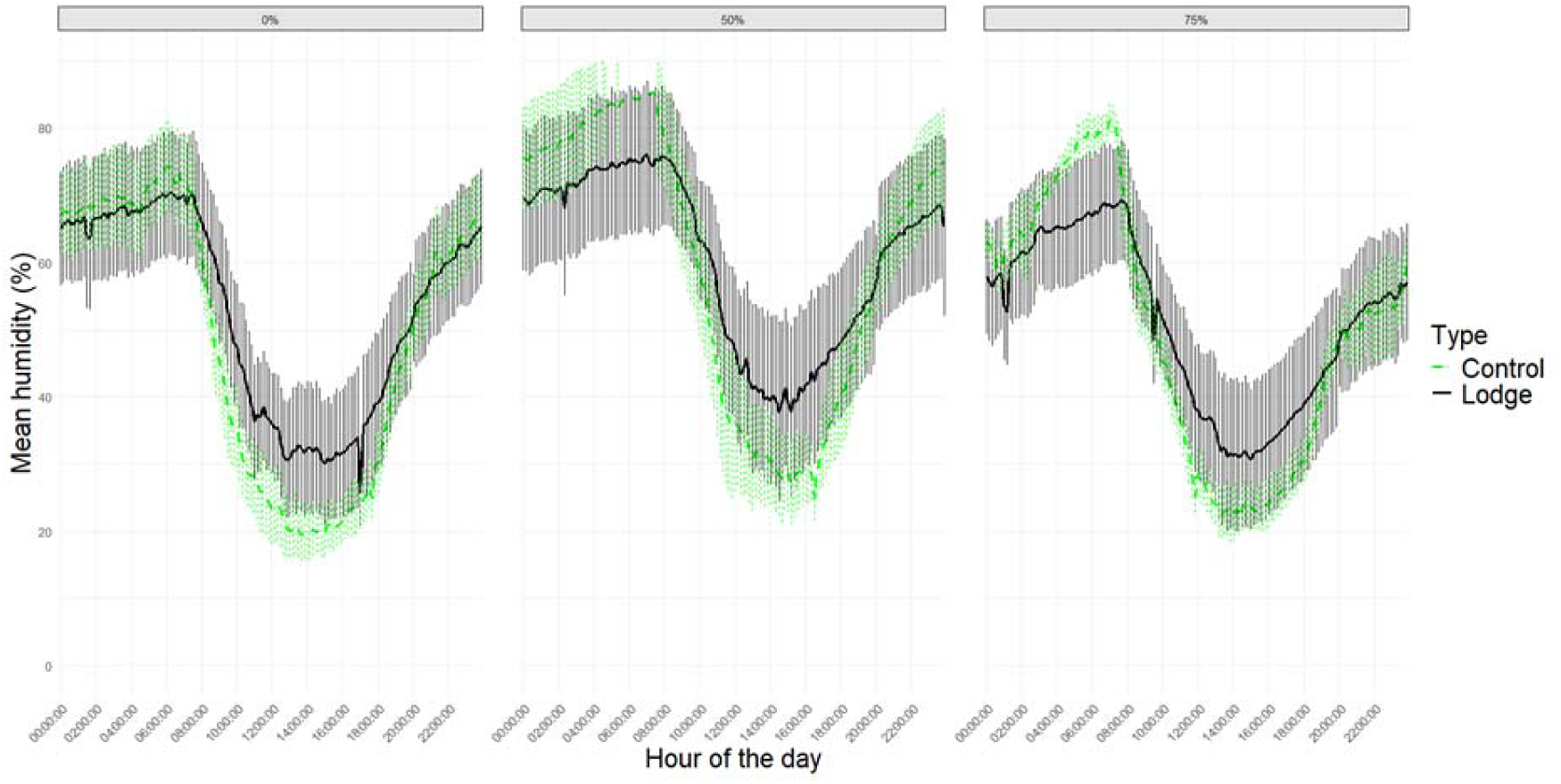
Relative humidity inside lodges depending on level of deconstruction, and ambient humidity.

## Discussion

Bush Karoo rat lodges are a striking example of animal architecture, constructed of up to 25 000 sticks and several kg of clay, faeces and sand, reaching weights of up to 48kg. Built by an animal weighing only around 100g, these impressive lodges display structural complexity in both externally and internally. External platforms provide basking sites. Internal microclimate conditions were closer to the thermoneutral zone of the bush Karoo rats than ambient temperature, and exhibited higher relative humidity. Together, these attributes likely reduce costs of thermoregulation and water loss, supporting our interpretation of the lodges as extended phenotypes.

Lodges’ dimensions varied considerably, with diameters ranging from 30-240cm, circumference from 58cm to 138cm, and heights from 12 to 74cm. Ours is the first study to measure all lodges within a study area instead of selected lodges, providing a comprehensive assessment of population-level architectural variation. Similarly, for other lodge building rodents such as American pack rats and the Australian stick-nest rat, data on size variation are rare or absent, because studies typically focused on large and old lodges (Betancourt, Van Devender & Martin 1990).

Lodge size was influenced by the shrub species into which they were built, with higher growing *Lycium* shrubs supporting taller lodges than the shorter *Tetraena* shrubs. All lodge measurements were strongly correlated, with taller lodges also having a greater circumference and diameter, and more entrances and platforms. This suggests that future studies could use a single metric as a proxy for overall lodge size. Seasonally, lodges were taller in the dry season, probably due to ongoing building activity by the bush Karoo rats, but also had a slightly smaller circumferences This may reflect reduced plant growth in the dry season, which might have improved measurement accuracy, and a decline in fresh vegetation carried to the lodge edges for drying and later consumption, so called plantong (Sangweni *et al*. 2025).

Externally, lodges were composed largely of woody material, although other items such as stones, mammalian dung and anthropogenic debris were occasionally incorporated. Bush Karoo rats had collected thousands to tens of thousands of sticks to build lodges, weighing on average 15kg. Clay and rodent faeces were the main materials for constructing platforms, which were present at most lodges. Platforms predominantly faced eastwards, providing terraces for morning basking. Basking to warm up passively has been observed in several species of small mammals (Geiser & Drury 2003; Stannard, Fabian & Old 2015) including the sympatric striped mouse (Schradin *et al*. 2007), where it is known to significantly reduce energy expenditure (Scantlebury *et al*. 2010). While striped mice select spots for basking inside shrubs that are warmer than random spots in the shrubs (Zduniak, Pillay & Schradin 2019), our findings suggest that bush Karoo rats construct dedicated basking structures into their architecture, perhaps providing the first empirical example of structural features explicitly constructed for this purpose.

Internally, lodges contained tunnels up to 75cm long, sleeping chambers, latrines and food chambers. Many tunnels appeared to have been blocked off with thorny branches. This raises the possibility that these internal barriers may impede predators, particularly snakes. Future studies should test whether these thorny barriers are in fact actively placed (and removed) by bush Karoo rats.

Ours is the first study to describe food chambers in addition to sleeping chambers and latrines, in bush Karoo rat lodges. These contained fresh food plants but lacked nest material and accumulated faeces. Food chambers likely allow bush Karoo rats to feed within the safety of the lodge while benefiting from more stable environmental conditions. Food chambers were as common than latrines, suggesting that they are functional components of lodges rather than incidental features. While Vermeulen and Nel (1988) reported two sleeping chambers per lodge (N=9), usually on different levels and near a latrine, our endoscope data did not provide reliable estimates of chamber numbers, because we often could not determine whether a tunnel ended or simply curved out of view. Thus, the number of internal features per lodge remains uncertain. Our lodge deconstruction also yielded limited information because we focused on abandoned lodges. Consequently, most internal structures were degraded, and few chambers and nests were found. This indicates that, beyond construction, lodge maintenance is a major time and energy investment for bush Karoo rats.

Lodges were clearly anchored within shrubs, using existing branches for stability into which collected sticks were placed. However, no consistent construction strategy using this pre-existing structure was evident. Most interesting, we found a ceiling-like structure at the top of either layer 1 and sometimes layer 2 and 3, composed of a hard mixture of dirt, faeces and sand. This is the first report of such a structure in bush Karoo rat lodges. The structure resembled human-made concrete, with faeces seemingly acting as the binding agent in place of cement, holding together sand and other materials into a cohesive mass. This is similiar amberat, formed from crystallized urine, which makes pack rat middens very hard and durable (Betancourt, Van Devender & Martin 1990). The ceiling layer may insulate the inner chambers from the external environment.

Bush Karoo rat lodges buffer environmental extremes. Compared to ambient conditions, internal temperatures were cooler during the heat of the day and warmer at night and relative humidity was consistently higher, as also reported previously (du Plessis, Kerley & Winter 1992; Jackson *et al*. 2002). The temperature differences averaged about 2°C and was similar or even higher than reported in Jackson *et al*. (2002). This would have allowed the bush Karoo rats to spend more time within their thermoneutral zone, thereby reducing energetic costs of thermoregulation. Relative humidity inside lodges was higher than outside during dry days, further improving the microclimate and reducing physiological costs (Gerson *et al*. 2014). Taken together, our study confirms that lodges provide a favorable microclimate, acting as a thermal refuge that reduces energetic costs in a harsh environment.

Although the favourable microclimate of bush Karoo rat lodges has been shown previously, so far nobody studied the relationship between microclimate and lodge size. We found that larger lodges provided a better microclimate. By experimentally reducing lodge size, we demonstrated that thermal buffering against heat declined with decreasing lodge size. Lodge size reduction led to lower humidity in the lodges and gradually increased internal temperature during the hottest hour of the day, although there was no effect of cold night temperatures, possibly because the study was conducted in summer, when night-time temperatures remained mild. Over a full 24-hour cycle, thermal conditions deviated further from the thermoneutral zone of the bush Karoo rats as lodge size decreased. Crucially, our experimental reduction of lodge size confirms that lodge size directly influences the microclimatic buffering.

Bush Karoo rat lodges are a striking example of how animals shape and modify their environment. Like many other rodents, they build warm nests of soft grass, but unlike most other rodents, they do not house these nests within burrows, but inside immense stick lodges that are up to 480 times the size of their builder. Compared to nests built by some fish (Stiassny & Meyer 1999; Östlund-Nilsson 2000) and most birds (Perez, Manica & Medina 2023), these lodges are remarkably large. For mammals, this is among the comparatively largest forms of animal architecture, perhaps only surpassed by beaver lodges (Baker & Hill 2003) as well as human buildings. Relatively (to builder) larger architectural structures are typically constructed only by social insects, and these result from the collective effort of large to very large colonies (Hölldobler & Wilson 2009), In contrast, the bush Karoo rat is a solitary species. Interestingly, these lodges are not built and maintained by a single individual but is a consequence of the accumulative efforts of several generations of individuals that use these lodges consecutively (Schradin & Makuya, pers. observ.). Constructing these lodges has obvious costs, distributed across generations, and our deconstruction of abandoned lodges with decayed internal structure indicates significant maintenance costs for every individual.

From an evolutionary perspective, the costs of lodge construction and maintenance must be outweighed by individual benefits. Functional benefits include predator avoidance, basking opportunities, thermal and hydric buffering and food storage. While no single function may explain the evolution of such structures, their combined value likely offsets construction and maintenance costs. Whether the selective pressure was primarily climatic, trophic or predatory remains unresolved. Future experiments could disentangle these roles, particularly by quantifying fitness consequences across lodge sizes and conditions. Lodges likely serve multiple, co-evolved functions, and represent an extended phenotype that increases the fitness of its builders.

Bush Karoo rat lodges meet all five criteria of extended phenotypes (Dawkins 1982; Laidre 2021). They (1) are external structures, extending the builder’s body, (2) are functional, (3) predicted to increase fitness, (4) result from behavioural activity, and (5) have properties predicted by the genotype. While the genetic predictions for lodge traits are untested, several lines of evidence suggest genetic determination in the general architecture. First, this species uses stick lodges consistently across its geographic range of several hundred kilometers in different climate and vegetation zones (Vermeulen & Nel 1988; Brown & Willan 1991; du Plessis, Kerley & Winter 1992; Kerley 1992; Jackson *et al*. 2002), suggesting a stereotyped behavioural template. Second, close relatives in the same genus do not construct such lodges but instead use burrows (Willan & Meester 1989; Hinze, Pillay & Grab 2006), indicating evolutionary divergence in nesting behaviour. Finally, bush Karoo rats are raised alone by their mother and often become independent before they start building lodges (Wolhuter *et al*. 2021), making social learning of lodge building unlikely. Therefore, the architecture of bush Karoo rat lodges is likely genetically encoded and constitutes an extended phenotype. The extended phenotype concept has historically been applied mostly to conspicuous structures built by invertebrates, such as spider webs, termite mounds or ant cemeteries (Nicholson 1963). In vertebrates, only a few examples have been suggested, including bowerbird constructions and stickleback nests (Laidre 2021). We consider bush Karoo rat lodges as a compelling example of an extended phenotype in a solitary vertebrate.

## Conclusions

Here we first quantified and described bush Karoo rat lodges as an example of animal architecture, then compared conditions inside versus outside the architecture, before experimentally testing for its function, complying with the research strategy suggested by Laidre (2021). These lodges, which were several hundred times the size of the builder, had complex external and internal structure, offering a favourable thermal and humidity microclimate. These benefits correlated with initial lodge size and decreased when lodges were experimentally reduced in size. This example of animal architecture likely improves thermoregulation, allows for predator avoidance and food storage, and thus represents an extended phenotype. Moreover, lodges might have significant effects on sociality, for example allowing solitary living by buffering the costs of missing out on benefits of group-living (Makuya & Schradin 2024). Also, by building lodges, bush Karoo rats modify local environmental conditions as eco-system engineers (Coggan, Hayward & Gibb 2018), a topic for future research. In conclusion, bush Karoo rat lodges are among the largest animal architecture made by solitary mammals. Bush Karoo rat lodges are complex, improve thermoregulation, allow for predator avoidance and food storage, and are actively maintained, represent a very good example of an extended phenotype.

## Acknowledgments

This study was made possible by the administrative and technical support of the Succulent Karoo Research Station (registered South African NPO 122-134). We are grateful for the support of Goegap Nature Reserve and its staff as well as the Northern Cape Department of Environment & Nature Conservation. This study was supported by the CNRS and the University of the Witwatersrand. This study is part of the long-term Studies in Ecology and Evolution (SEE-Life) programme of the CNRS. This project has received financial support from the CNRS through the MITI interdisciplinary programs through its exploratory research program. Further funding was provided by the French National Research Agency (ANR) for the project COBESOLI.

## Data Availability

Data and code will be made available on indores after publication (Schradin *et al*. 2025). Its available for review under the following link: https://data.indores.fr:443/privateurl.xhtml?token=c3bb7013-9a3a-4d69-8f0e-0381c01401da **Declaration of Interest**

The authors declare that they have no conflicting interests.

**Supplementary material** associated with this article will be made available in the online version.

