## supplemental tables and figures for "Thermal Refuges as Extended Phenotypes: Lodge Construction by Bush Karoo Rats"

**ESM**

**Model 1: Height ~ Diameter + Shrub + Season + (1|Lodge)**

- **Fixed Effects:**
  - Intercept = 28.21 ± 2.81, t(df = 179.31) = 10.03, p < 0.001
  - Diameter = 0.18 ± 0.02, t(df = 203.39) = 8.66, p < 0.001
  - Shrub (Zygophyllum) = –9.68 ± 1.90, t(df = 143.90) = –5.08, p < 0.001
  - Season (Moist) = –2.71 ± 1.03, t(df = 111.91) = –2.65, p = 0.0093
- **Random Effect (Lodge ID):**
  - Variance = 33.49; Residual = 54.52
- **Model Fit:** REML = 1678.7

**Model 2: Height ~ Circumference + Shrub + Season + (1|Lodge)**

- **Fixed Effects:**
  - Intercept = 22.53 ± 2.96, t(df = 161.56) = 7.60, p < 0.001
  - Circumference = 0.07 ± 0.01, t(df = 167.77) = 9.96, p < 0.001
  - Shrub (Zygophyllum) = –7.32 ± 1.77, t(df = 139.17) = –4.13, p < 0.001
  - Season (Moist) = –5.14 ± 1.11, t(df = 128.22) = –4.61, p < 0.001
- **Random Effect (Lodge ID):**
  - Variance = 19.25; Residual = 61.41
- **Model Fit:** REML = 1669.7

**Model 3: Height ~ Number of Entrances + Shrub + Season + (1|Lodge)**

- **Fixed Effects:**
  - Intercept = 42.77 ± 2.54, t(df = 195.91) = 16.86, p < 0.001
  - Number of Entrances = 1.38 ± 0.47, t(df = 218.42) = 2.93, p = 0.0038
  - Shrub (Zygophyllum) = –7.37 ± 2.33, t(df = 156.96) = –3.16, p = 0.0019
  - Season (Moist) = –2.90 ± 1.15, t(df = 107.53) = –2.53, p = 0.0128
- **Random Effect (Lodge ID):**
  - Variance = 53.03; Residual = 63.61
- **Model Fit:** REML = 1692.5

**Model 4: Height ~ Number of Platforms + Shrub + Season + (1|Lodge)**

- **Fixed Effects:**
  - Intercept = 39.68 ± 2.75, t(df = 182.71) = 14.43, p < 0.001
  - Number of Platforms = 1.79 ± 0.45, t(df = 214.34) = 4.00, p < 0.001
  - Shrub (Zygophyllum) = –4.83 ± 2.47, t(df = 157.34) = –1.96, p = 0.0520 (marginal)
  - Season (Moist) = –2.05 ± 1.12, t(df = 111.20) = –1.83, p = 0.0701 (marginal)
- **Random Effect (Lodge ID):**
  - Variance = 50.27; Residual = 62.74
- **Model Fit:** REML = 1679.2

**Table S1** Descriptive statistics of internal characteristics of 89 lodges where we could access the tunnels to insert the endoscope camera.

| **Chambers** | Mean ± SD (range; N=total number) for 89 lodges | **Comments** |
| --- | --- | --- |
| Number of Chambers (total) | 0.49 ± 0.74 (0-3; N=44) |  |
| Number of nest chambers (total) | 1.06 ± 0.24 (0-2; N= 18) |  |
| Number of latrine chambers (total) | 1.2 ± 0.42 (0-2; N=12) |  |
| Number food storage chambers (total) | 1.15 ± 0.55 (0--3; N=13) |  |
| Number food storage + latrine chambers (total= | 1 ± NA (0-1; N=1) | Both feces and food were seen in the chamber |
| **Tunnels** |  |  |
| Number of tunnels | 3.07 ± 1.47 (0-7; N= 271) |  |
| Length of tunnels | 34.11 ± 13.8cm (10-75; N=271) |  |
| Number of tunnels leading to: inside | 1.47 ± 1.22 (0-5; N= 131) | No chamber or any special lodge structure seen at end. |
| Number of tunnels leading to: nest | 0.16 ± 0.37 (0-1; N=14) | C or U shape structure, lines with grass material |
| Number of tunnels leading to: latrine chamber | 0.16 ± 0.42 (0-2; N=14) | C or U shape structure with lots of feces |
| Number of tunnels leading to: food chamber | 0.10 ± 0.30 (0-1; N=9) | C or U shape structure with lots of fresh plant material being stored |
| Number of tunnels leading to: latrine + food chamber | 0.02 ± 0.19 (0-1; N=2) | Both fresh plant material and fecal are stored in the same structure |
| Number of tunnels leading to: platform | 0.24 ± 0.54 (0-2; N=21) | The platform is the special flat structure for many activities, is the veranda of the lodge |
| Number of tunnels leading to: tunnel | 0.08 ± 0.38 (0-3; N=7) | The main tunnel might lead to another tunnel inside the same lodge |
| Number of tunnels leading to underground | 0.12 ± 0.39 (0-2; N=11) | The tunnel can lead to the base or ground and continue to straight to burrows. |
| Number of tunnels leading to ground | 0.30 ± 0.64 (0-4; N=27) | Ground is the base of the lodge, where lodge was built |
| Number of tunnels leading to: outside | 0.35 ± 0.60  (0-3; N=31) | The lodge tunnel can lead to inside then outside of the lodge |

**Photos internal structure**

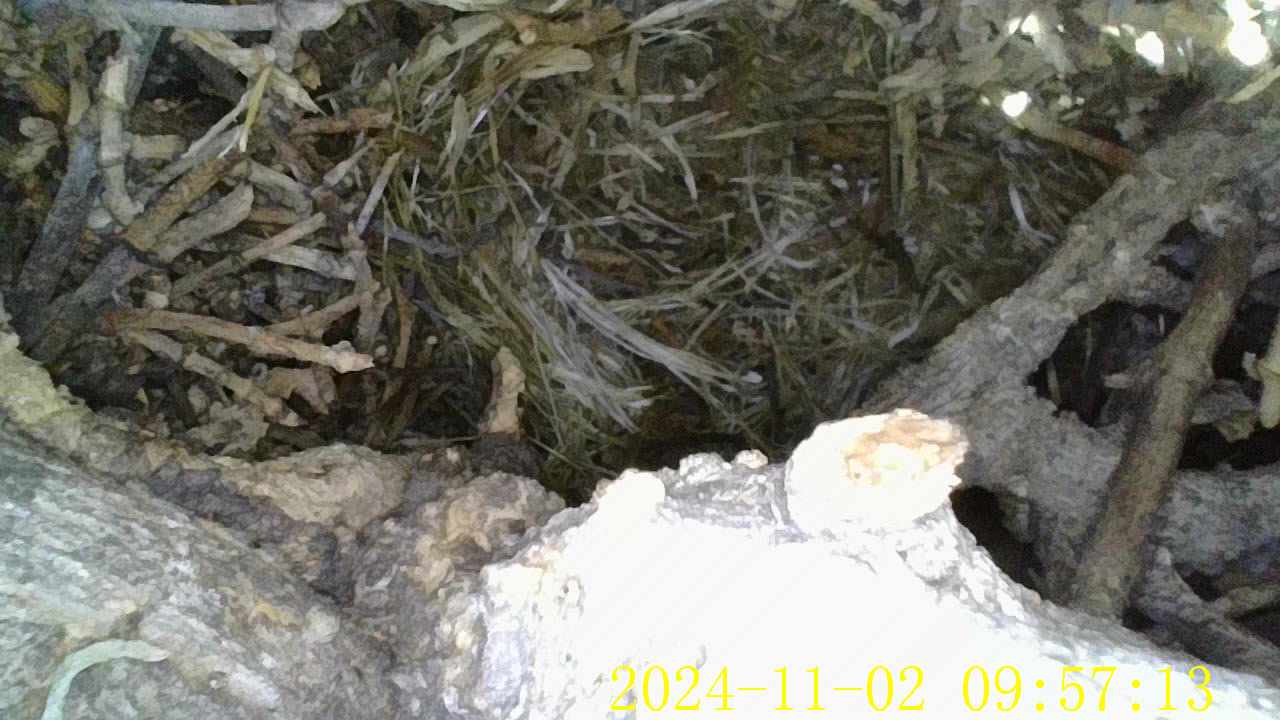

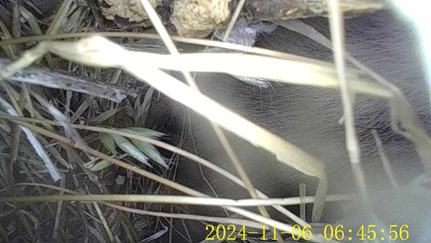

**Figure S1: Nesting chambers**

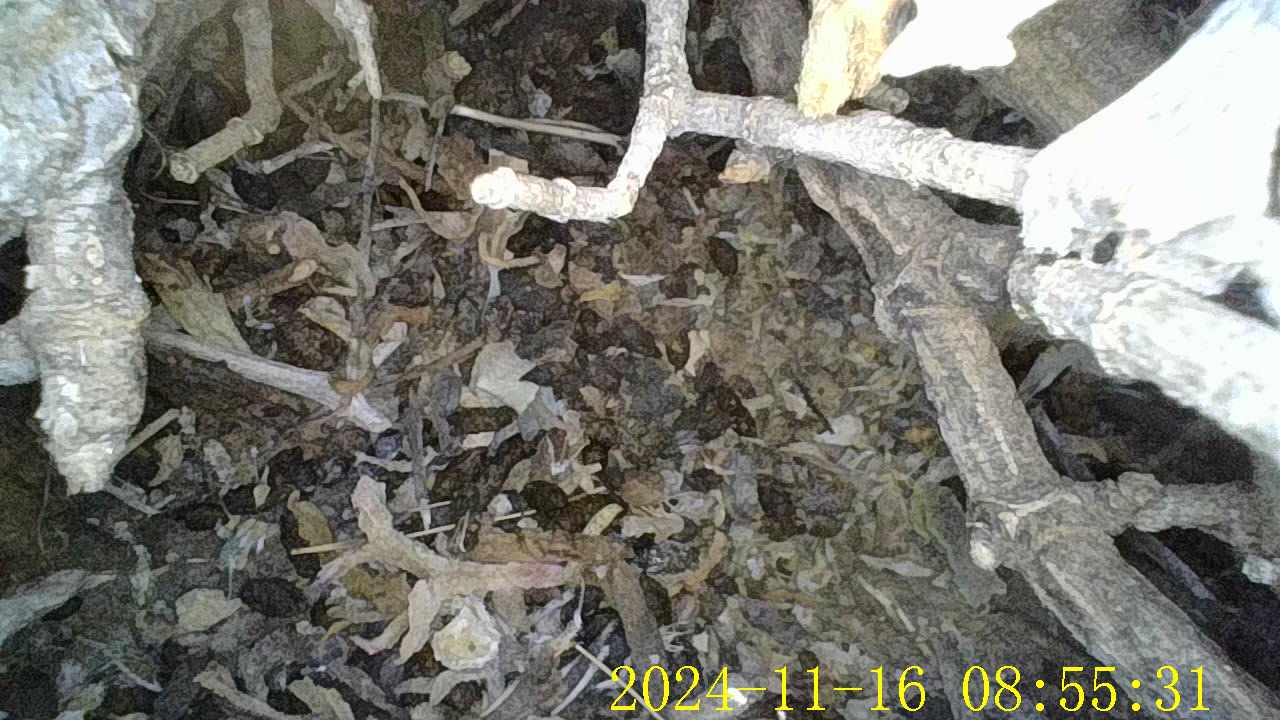

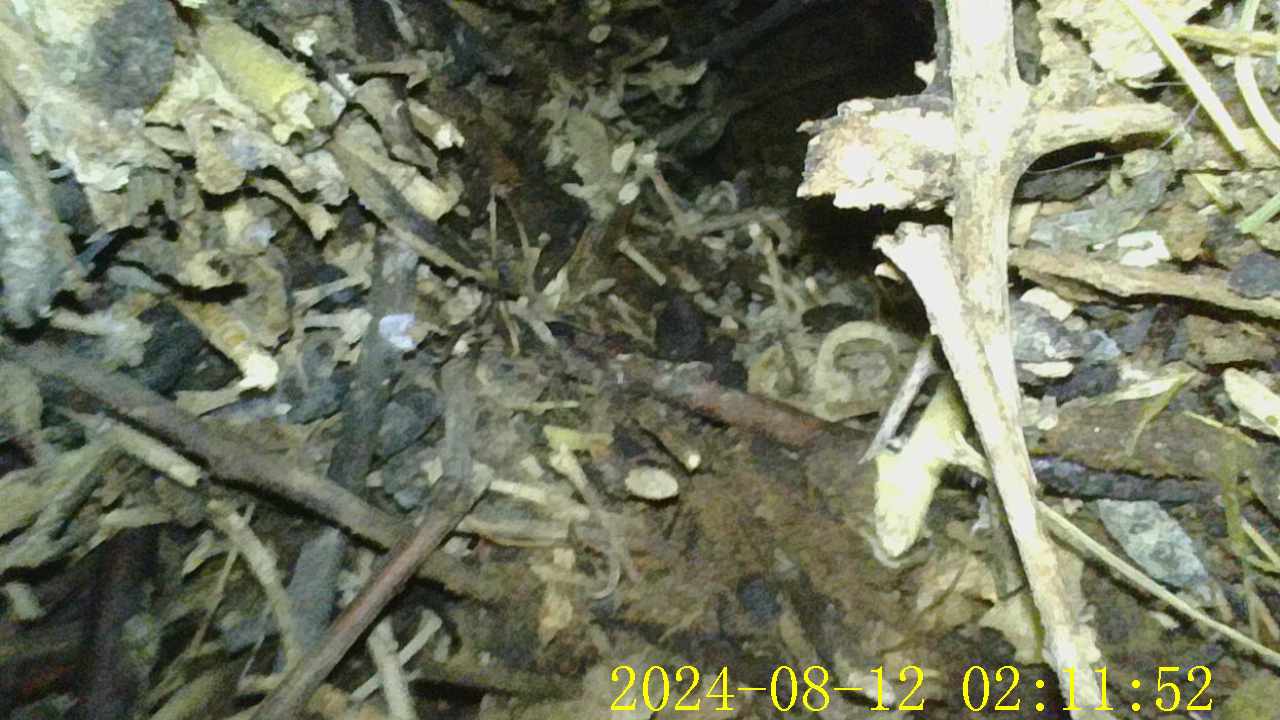

**Figure S2: latrines**

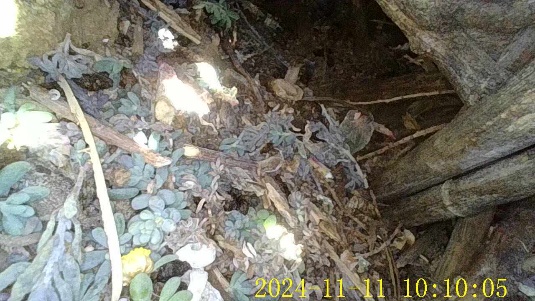

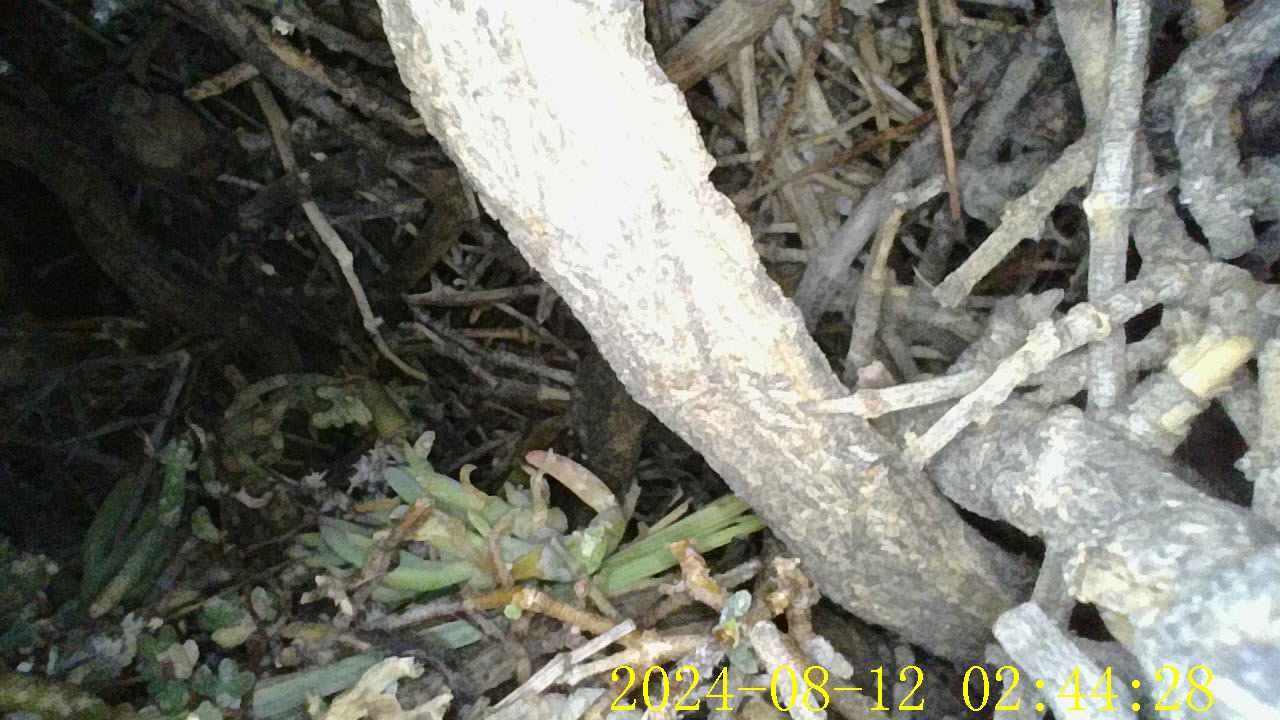

**Figure S3: food chambers**

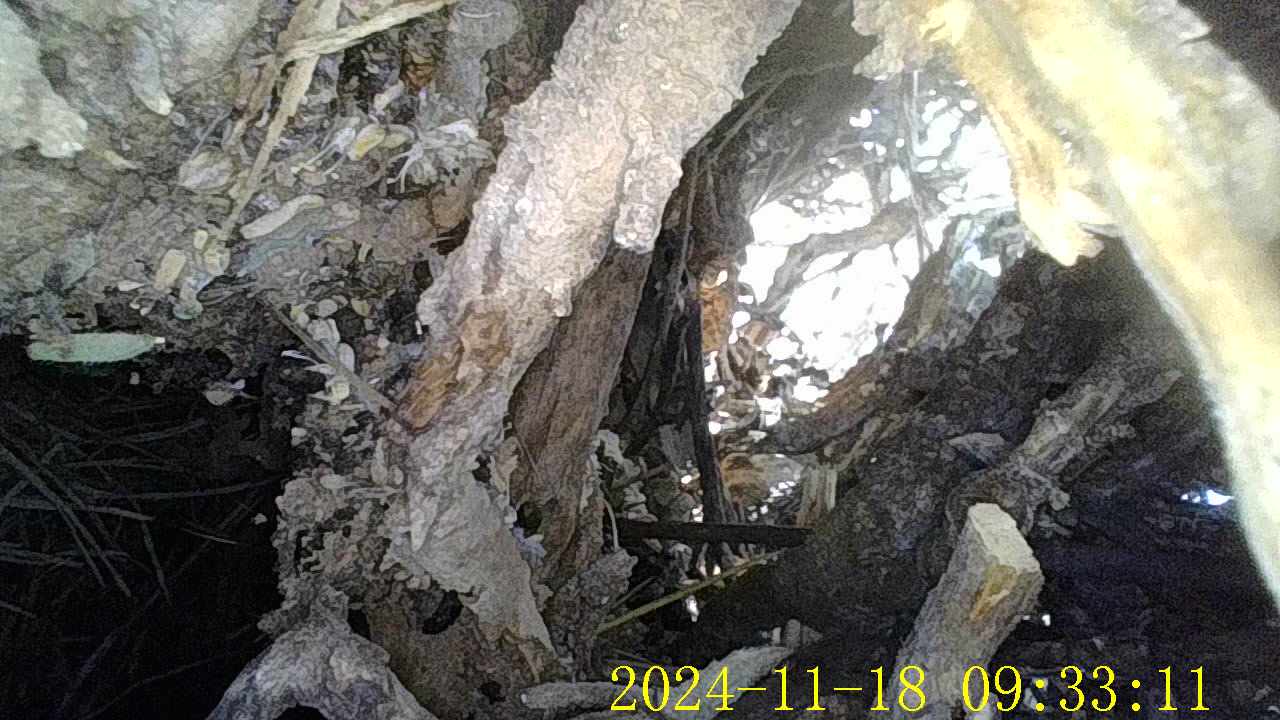

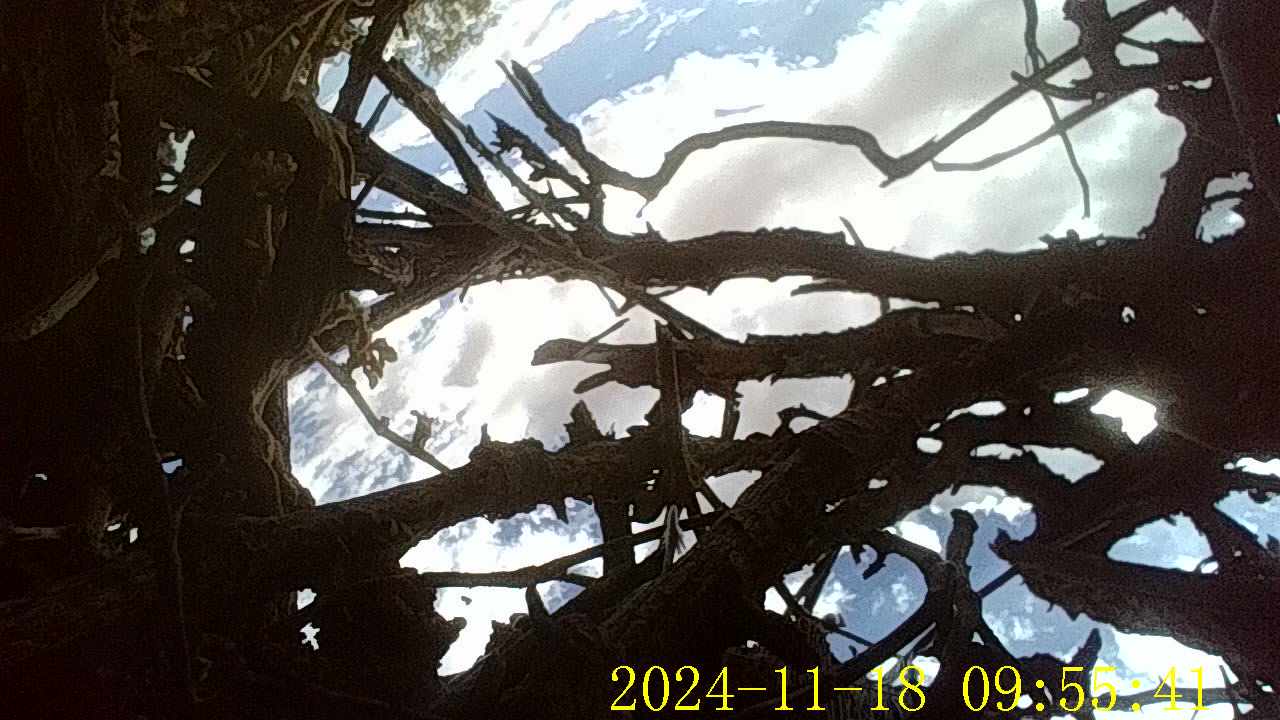

**Figure S4: tunnels leading to the outside (2 photos)**

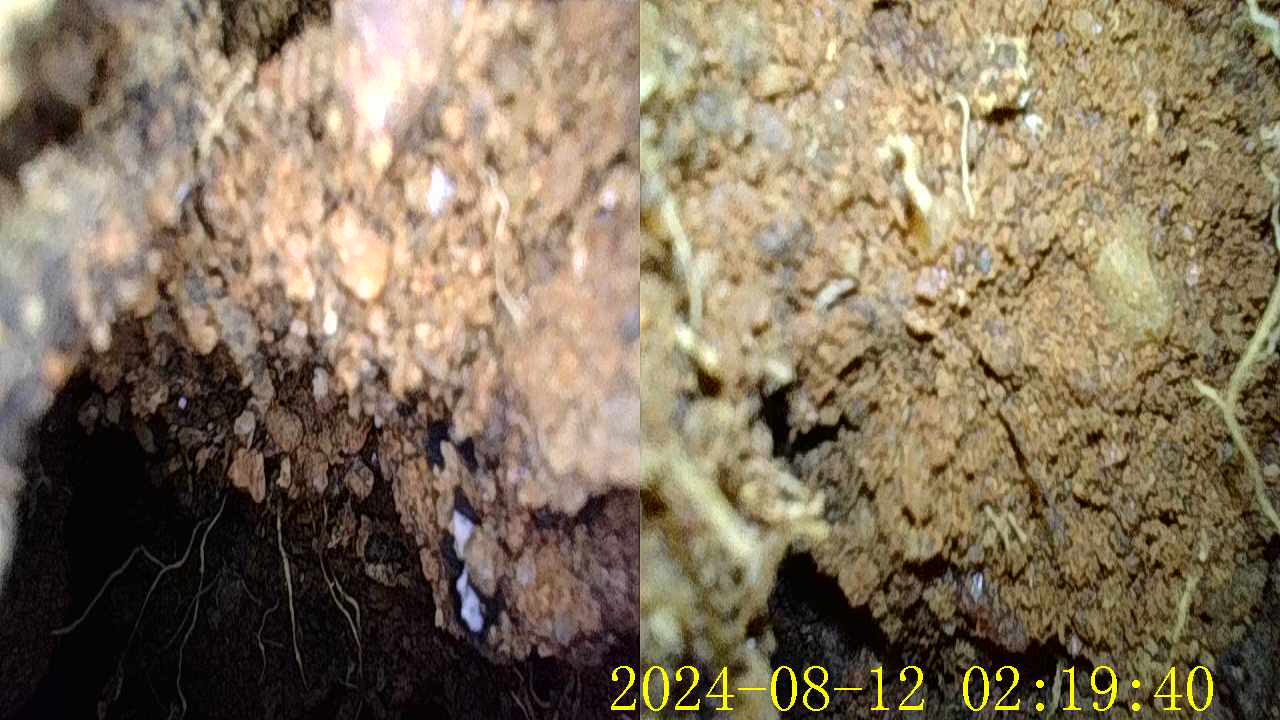

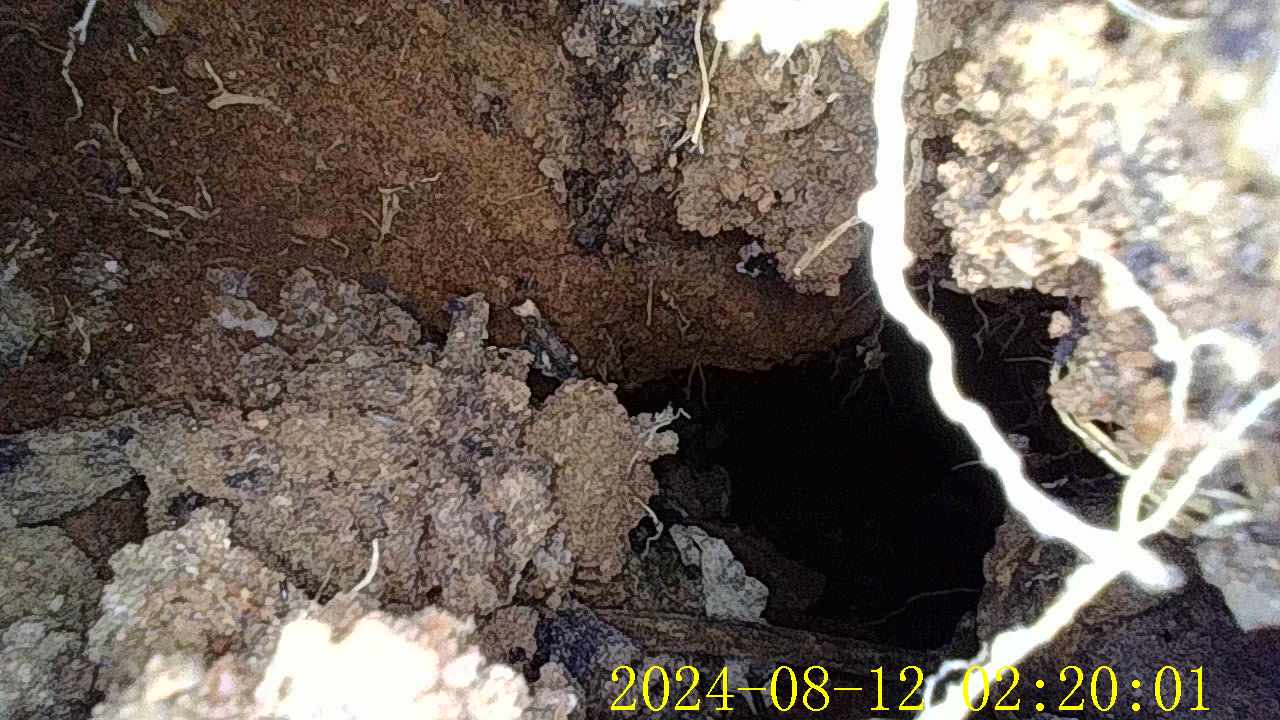

**Figure S5: the underground floor of the lodge**

**Table S2**

Descriptive statistics of sticks from different layers of the eight deconstructed lodges, all of which had at least 4 layers.

|  | Layer 1 | | Layer 2 | | Layer 3 | | Layer 4 | | Layer 5 (n= 5) | |
| --- | --- | --- | --- | --- | --- | --- | --- | --- | --- | --- |
|  | Number | Mass (g) | Number | Mass (g) | Number | Mass (g) | Number | Mass (g) | Number | Mass (g) |
| Sticks 1-5cm | 4083 ± 3189 | 849 ± 876 | 2502 ± 2195 | 402 ± 386 | 1004 ± 1333 | 129 ± 157 | 170 ± 253 | 23 ± 41 | 9 ± 13 | 1 ± 1 |
| Sticks >5-10cm | 1866 ± 1481 | 837 ± 858 | 1667 ± 1000 | 709 ± 610 | 1253 ± 1244 | 548 ± 522 | 260 ± 295 | 195 ± 239 | 32 ± 26 | 51 ± 42 |
| Sticks >10-15cm | 254 ± 178 | 430 ± 475 | 358 ± 128 | 417 ± 352 | 398 ± 259 | 544 ± 541 | 154 ± 89 | 307 ± 310 | 51 ± 37 | 133 ± 159 |
| Sticks >15-20cm | 47 ± 41 | 192 ± 213 | 76 ± 24 | 173 ± 142 | 125 ± 57 | 284 ± 262 | 83 ± 48 | 267 ± 315 | 27 ± 12 | 89 ± 86 |
| Sticks >20cm | 22 ± 24 | 193 ± 256 | 31 ± 13 | 148 ± 127 | 53 ± 41 | 268 ± 334 | 45 ± 28 | 255 ± 350 | 26 ± 16 | 182 ± 159 |
| Total sticks | 6272 ± 4801 | 10069 ± 8937 | 4634 ± 3256 | 1850 ± 1578 | 2832 ± 2795 | 1773 ± 1640 | 711 ± 674 | 1047 ± 1226 | 90 ± 94 | 326 ± 415 |
| Stones |  | 13±29 |  | 25 ± 50 |  | 17 ± 24 |  | 3 ± 7 |  | 0 |
| Dung large ungulates |  | 3±5 |  | 1 ± 1 |  | 15 ± 35 |  | 11 ± 31 |  | 0 |
| Dung Jackals |  | 3 ± 5 |  | 0 ± 1 |  | 0 ± 1 |  | 0 |  | 0 |
| Mix of dirt and feces (g) |  | 7611 ± 7228 |  | 4065 ± 3946 |  | 684 ± 1279 |  | 37 ± 79 |  | 0 |
| **Total mass of lodge material** |  | **17704 ± 16070** |  | **3910 ± 3907** |  | **2230 ± 1921** |  | **1084 ± 1218** |  | **285 ± 401** |
